# Temperature-dependent polar lignification of a seed coat suberin layer promoting dormancy in Arabidopsis thaliana

**DOI:** 10.1101/2024.07.09.602674

**Authors:** Lena Hyvärinen, Christelle Fuchs, Anne Utz-Pugin, Kay Gully, Christian Megies, Julia Holbein, Mayumi Iwasaki, Lara Demonsais, Maria Beatriz Capitão, Marie Barberon, Rochus Franke, Christiane Nawrath, Sylvain Loubéry, Luis Lopez-Molina

## Abstract

The seed is a landmark plant adaptation where the embryo is sheltered by a protective seed coat to facilitate dispersion. In Arabidopsis, the seed coat, derived from ovular integuments, plays a critical role in maintaining dormancy, ensuring germination occurs during a favorable season. Dormancy is enhanced by cold temperatures during seed development by affecting seed coat permeability through changes in apoplastic barriers. However, the localization and composition of these apoplastic barriers are poorly understood. This study identifies and investigates a polar barrier in the seed coat’s outer integument (oi1) cells. We present histological, biochemical, and genetic evidence showing that cold promotes polar seed coat lignification of the outer integument 1 (oi1) cells and suberization throughout the entire oi1 cell boundary. The polar oi1 barrier is regulated by the transcription factors MYB107 and MYB9. MYB107, in particular, is crucial for the lignified polar oi1 barrier formation under cold temperatures. The absence of the oi1 barrier in mutant seeds correlates with increased permeability and reduced dormancy. Our findings elucidate how temperature-induced modifications in seed coat composition regulate dormancy, highlighting the roles of suberin and lignin in this process.

**Significance statement:** Our study uncovers how cold temperatures during seed development in the mother plant influence seed dormancy through apoplastic modifications in the *Arabidopsis thaliana* seed coat. We identified a polar lignin barrier in the outer integument 1 (oi1) cells, which are also suberized. Lignification and suberization are regulated by transcription factors MYB107 and MYB9. Cold promotes lignification and suberization of oi1 cells through MYB107, thus creating a “memory” that reduces seed permeability and strengthens dormancy. Mutants defective in the oi1 barrier exhibit lower dormancy, highlighting the adaptive importance of this barrier. These findings advance our understanding of temperature-induced seed coat adaptations and their agricultural implications, particularly in the context of climate change, offering valuable insights for improving crop resilience and yield.

## Introduction

The seed is a major terrestrial plant innovation, encapsulating the plant embryo within a protective seed coat to facilitate dispersion (1, 2). The mature Arabidopsis seed comprises an embryo encased by a single cell layer of endosperm, which is itself enclosed by the seed coat–a dead maternal tissue arising from ovular integuments. In Arabidopsis these integuments consist of five cell layers: three cell layers of inner integuments (ii1, ii1’ and ii2) and two cell layers of outer integuments (oi1 and oi2). When newly produced, mature dry seeds are dormant, a trait whereby seed germination is blocked even under favorable conditions, aiding seed dispersion and ensuring seedling establishment during favorable seasons. Dormancy is lost over time and the duration of dry after-ripening time to become non-dormant defines seed dormancy levels. Temperature is a major seasonal cue and cold temperatures (10-15°C) during seed development increase dormancy in many plants, including Arabidopsis (3–5). Hence, plants form anatomically fixed structures irrespective of environmental fluctuations, but their seed progeny can “remember” past maternal cold temperatures, resulting in increased dormancy.

The mechanisms sustaining dormancy release in dry seeds are not well understood but may involve diffusion of atmospheric oxygen within the seeds which triggers oxidative events that release dormancy (6). In Arabidopsis, the seed coat promotes dormancy, and studies have shown that cold enhances dormancy by modifying seed coat apoplastic barriers, serving as a memory of past temperatures. However, the specific identity and location of these modifications remains to be fully understood. The best characterized example is that of *transparent testa* (*tt*) mutant seeds, which have a more permeable seed coat and lack dormancy (7). *tt* seed coats lack flavonoids, which are antioxidants and are the building blocks of tannins accumulating in the ii1 cell walls, forming a reticulate structure surrounding the seed’s living tissues (8, 9). Cold during seed development increases levels of procyanidins that polymerize to form condensed tannins and *tt* seeds produced under cold conditions also have low dormancy levels (10, 11).

Available evidence suggests that temperature affects additional, yet unknown, seed coat apoplastic barriers regulating dormancy. Suberin is an aliphatic polyester accumulating between the plasma membrane and the inner cell wall. Glycerol-3-phosphate O-acyltransferase 5 (GPAT5) is involved in suberin synthesis (12). Suberin was proposed to be produced in the oi1 layer and Molina et al. suggested that *GPAT5* is expressed in oi1 cells (13–15). Furthermore, presence of lamellae in transmission electron microscopy (TEM) images, attributed to suberin depositions, were reported in the inner part of oi1 cells in seeds produced under warm temperatures (15, 16). Furthermore, MYeloBlastosis (MYB) family transcription factors (TFs) MYB107 and MYB9 promote suberin biosynthetic gene expression in seeds and *myb107* mutant seeds have disordered lamellae in the inner part of oi1 cells (16, 17). However, the presence of suberin in oi1 cells was not further investigated genetically and it remains unclear whether lamellae are present throughout the surface of oi1 cells, whether they encircle the seed’s living tissues or whether they are only present in specific seed coat locations (Chang-Jun Liu, personal communication) (15, 16). In addition, no major dormancy phenotypes in *myb107* or *myb9* mutant seeds were reported (16, 17). Furthermore, whether *MYB107* and *MYB9* regulate polyester or other depositions (such as lignin, see below) in response to cold temperatures to promote dormancy was not investigated. Indeed, cold temperatures affect the overall polyester content of seeds. When seeds are produced under cold conditions, both *awake1* (*awe1*) seeds, which are deficient in the ATP-binding Cassette transporter of class G ABCG20 that promotes suberin deposition, and *gpat5* seeds exhibit low dormancy (15, 18). Hence, it remains to be clarified which suberin-containing seed coat apoplastic barriers promote dormancy in response to cold.

Like tannin, lignin is a phenylpropanoid-based polymer that is assembled from monolignols (hydroxycinnamyl-alcohols) and is predominantly deposited in the cell wall conferring impermeability (19). Interestingly, MYB107 was shown to regulate phenylpropanoids and lignin biosynthesis gene expression in seeds; however, whether lignin is present in the oi1 layer was not shown (17). Histological evidence suggests presence of lignin in the seed abscission zone (20). *TT10/AtLAC15*, encoding a laccase-like polyphenol oxidase, is specifically expressed in ii1 and oi1 cells and is involved in the polymerization of seed flavonoids and monolignols (21, 22). *tt10* mutant seeds produced under warm temperatures have both lower lignin and tannin content, making it difficult to separate their roles in seed physiology (21). Hence, it is unclear whether lignin is present in the seed coat and whether it could be involved in promoting dormancy.

In conclusion, evidence strongly suggests that temperature during seed development alters the seed coat’s apoplastic barriers to promote dormancy. However, the localization, composition, and temperature-induced changes of these barriers remain poorly understood.

We present histological, biochemical and genetic evidence that oi1 cells form a polar apoplastic barrier with lignin on their outer sides and suberin lining the entire cell boundary. Lignification is weak in seeds produced under warm temperatures but reinforced under cold temperatures. *MYB107* and *MYB9* are specifically expressed in oi1 cells and promote oi1 barrier formation during the mature green stage with MYB107 playing a key role under cold temperatures. The absence of this barrier correlates with high seed coat permeability and low dormancy in both seeds produced under warm or cold temperatures. Thus, we identified a seed coat apoplastic barrier containing suberin and lignin, regulated by cold to promote seed dormancy.

## Results

### Identification of a polar oi1 barrier reinforced by cold during seed development

We sought to identify seed coat apoplastic barriers being formed or modified by cold during seed development. WT (Col-0) Arabidopsis plants were grown at 22°C or transferred at bolting to 13°C. Mature dry seeds produced at 22°C and 13°C are referred to as ‘Warm seeds’ and ‘Cold seeds’, respectively. Histological sections of Warm and Cold seeds were stained with Auramine O (AurO), a fluorescent dye staining cutin, suberin and lignin (9, 23). Strikingly, cold induced a strong linear periclinal signal between differentiated and dead oi1 and oi2 cells (hereafter ‘oi1 cells’ and ‘oi2 cells’) and surrounding the seed’s living tissue (Fig. 1A, S1A & S1B). In addition, this signal was interspersed with short anticlinal signals running between adjacent oi1 cells (Fig. 1A). A similar, but weaker, signal was present in Warm seeds, revealed by increased image contrast (Fig. 1A, S1A & S1B). These observations suggest the presence of a polar barrier revealed by strong outer periclinal and anticlinal AurO signals, referred to as the ‘polar oi1 barrier’ whose composition is markedly regulated by cold. Contrasted images showed the inner periclinal and inner anticlinal side of oi1 cells also stained by AurO, referred to as the ‘inner oi1 barrier,’ with increased intensity in Cold seeds although markedly weaker relative to that of the polar oi1 barrier signal (Fig. 1A). This indicates that cold also regulates the inner oi1 barrier composition, differing quantitatively or qualitatively from that of the polar oi1 barrier. No noticeable changes in the fluorescent signal from the ii1 cuticle associated with the endosperm were observed between Warm and Cold seeds (Figs.1A and S1C) (24).

**Figure 1:**
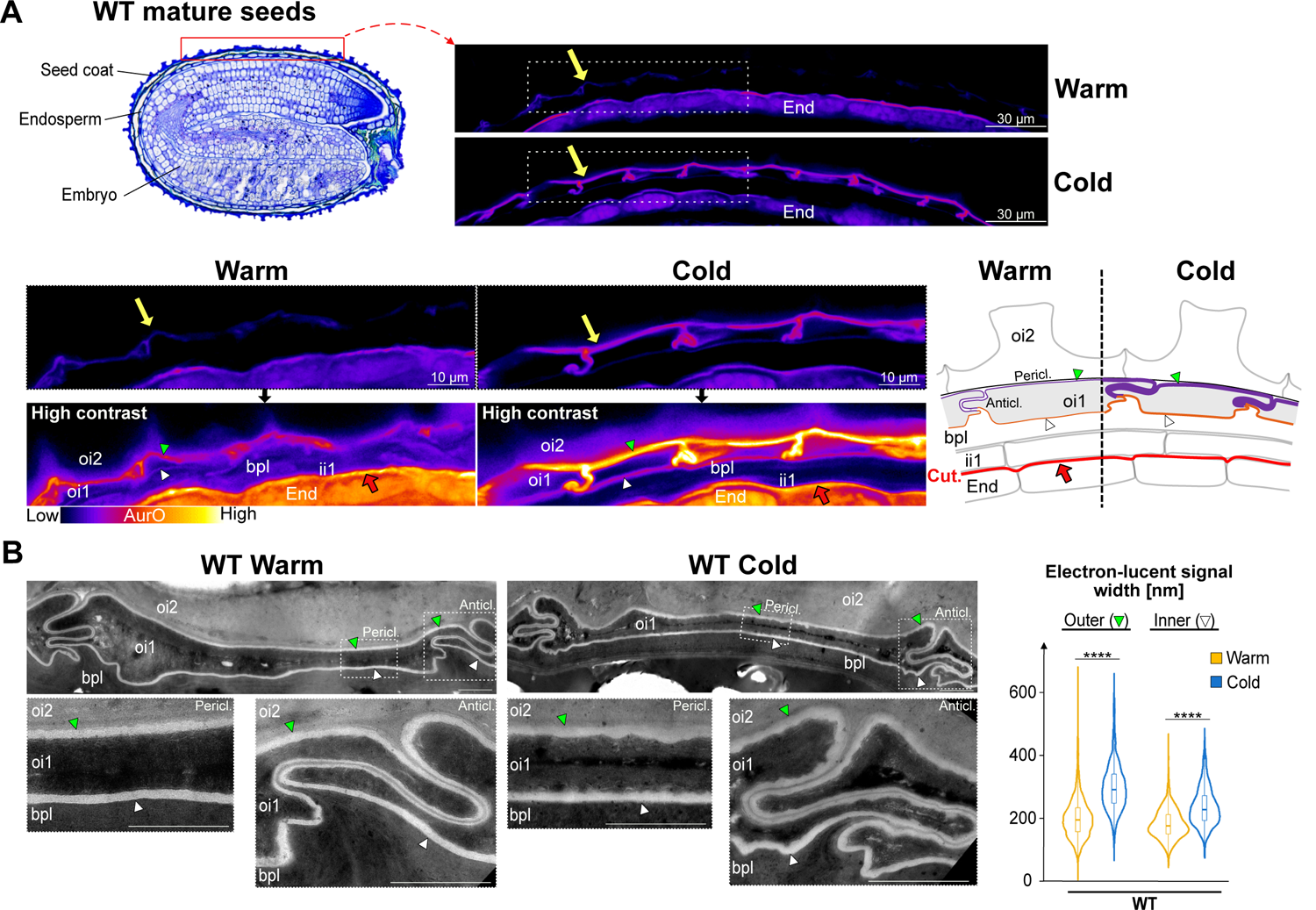
Identification of a polar oi1 barrier reinforced by cold during seed development. **A.** Top left: histological section of a WT mature seed stained with Toluidine Blue; the red rectangle indicates the region examined on the top right panel. Top right: AurO staining of WT mature seed sections showing the region indicated by the red rectangle in seeds that developed under warm (Warm) temperatures (22°C) or cold (Cold) temperatures (13°C). Yellow arrow indicates the AurO signal corresponding to the polar oi1 barrier. Bottom left: magnifications of the regions in top right panel delimited by a dashed white rectangle in WT Warm and Cold seeds; a higher contrast is also shown, as indicated. Bottom right: seed coat schematic depicting in Warm and Cold seeds the polar (violet line) and inner (brown line) oi1 barrier; the thick red line shows the ii1 cuticle (Cut.) associated with the endosperm. Green arrowhead, polar oi1 barrier; white arrowhead, inner oi1 barrier, red arrow indicates ii1 cuticle; Pericl.: periclinal; Anticl.: anticlinal. **B.** Top left: TEM micrographs showing the linear electron-lucent signal surrounding oi1 cells in WT Warm and Cold seeds. Bottom left: magnification of the anticlinal (Anticl.) and periclinal (Pericl.) regions in top left panel delimited by dashed white rectangles in WT Warm and Cold seeds as indicated. Green and white arrowheads indicate the outer and inner oi1 electron-lucent signal, respectively. Right: Violin plots of the outer and inner electron-lucent signal width, n=6 cells (3 seeds, 2 cells per seed), statistical analysis as assessed by Kruskal-Wallis test (****p< 0.0001). green and white arrowhead, outer and inner oi1 electron-lucent signal, respectively. oi2, outer integument 2; oi1, outer integument 1; bpl, brown pigmented layer (fusion of inner integument 1’ and ii2 layers); ii1, inner integument 1 layer; End., endosperm. Bar, 2µm.

Consistent with previous reports, transmission electron microscopy (TEM) revealed a smooth and linear electron-lucent signal surrounding oi1 cells in Warm seeds, which was previously attributed to suberin depositions (16) (Fig.1B). In Cold seeds, a similar electron-lucent signal was observed in oi1 cells although it was moderately thicker and more irregular relative to Warm seeds (Fig. 1B). Interestingly, in both WT Warm and Cold seeds, this signal had a prototypical shape, which to our best knowledge was not previously recognized: the outer anticlinal electron-lucent signals of two adjacent cells run closely to each other and form convolutions, while the inner anticlinal signals separate, creating a pointed hat-shaped structure (Fig. S1D). This pattern suggested the occurrence of a polarization in oi1 cells, consistent with the polar AurO signal (see discussion). Accordingly, the convoluted outer anticlinal signals, together with the outer periclinal signals, form a pattern like that of the polar oi1 barrier seen with AurO in Cold seeds, as can be appreciated in the contrasted images (Fig. 1A). In Warm seeds, no obvious polarity in either electron density or width was observed between the outer and inner signals (Fig. 1B). In Cold seeds, the width of the outer and inner signals increased relative to Warm seeds, suggesting that Cold promotes oi1 suberization (Fig. 1B). However, the outer periclinal signal became only moderately wider than the inner one (Fig. 1B). Hence, the TEM electron-lucent oi1 signal does not satisfactorily explain the occurrence of the polar AurO signal in Warm seeds and its marked increase in Cold relative to Warm seeds (Figs. 1A & S1B). Therefore, the composition of the polar oi1 barrier detected by AurO remains to be clarified (see below).

We aimed to identify when the polar oi1 barrier appears during seed development. Coen et al. used AurO to stain seeds up to the torpedo stage, i.e. 7 to 8 days after pollination (DAP), finding no signal in the oi1 cell layer, indicating that the polar oi1 barrier forms after this stage (25). We detected no AurO signal at 12 DAP (early mature green stage) but found an apolar signal around oi1 cells at 15 DAP (mid-mature green stage) and 18 DAP (late-mature green stage) (Fig. S1E). This indicates that polarization is completed at later stages. Similar results were observed in seeds developing at 13°C: no AurO signal during the walking stick stage, but detectable during the mature green stage (Fig. S1F).

### MYB107 is essential for polar oi1 barrier formation in seeds developing under cold temperatures

To characterize the polar oil barrier’s composition, we sought to identify TFs crucial for its formation under cold temperatures. In turn, mutants lacking these TFs may help pinpoint misregulated gene expression in seeds, offering insights into the barrier’s composition.

MYB9 and MYB107 were found to promote suberin and phenylpropanoid biosynthetic gene expression in whole seeds (16, 17). While WT, *myb9-1*, and *myb107-2* Warm seeds showed similar polar AurO signals (Fig. 2A), a weaker signal was detected in *myb9 myb107* Warm seeds (Fig. S2A), indicating that *MYB9* and *MYB107* play a redundant role in polar oi1 barrier formation in Warm seeds. In *myb9-1* Cold seeds, the polar oi1 barrier AurO signal increased compared to *myb9-1* Warm seeds, though less than in WT Cold seeds (Fig. 2A). Strikingly, the signal was absent in *myb107-2* Cold seeds (Fig. 2A), which was confirmed with the independent *myb107-1* mutant allele (Fig. S2B). Thus, MYB107 predominantly promotes polar barrier formation under cold temperatures, with MYB9 playing a lesser role. Furthermore, F1 Cold seeds obtained after pollinating *myb107-2* plants with WT pollen did not produce a polar oi1 barrier, unlike F1 Cold seeds arising from the reciprocal cross (Fig. S2C). Hence, MYB107 activity in the maternal seed coat plays a predominant role under cold temperatures to promote the formation of the polar oi1 barrier.

**Figure 2:**
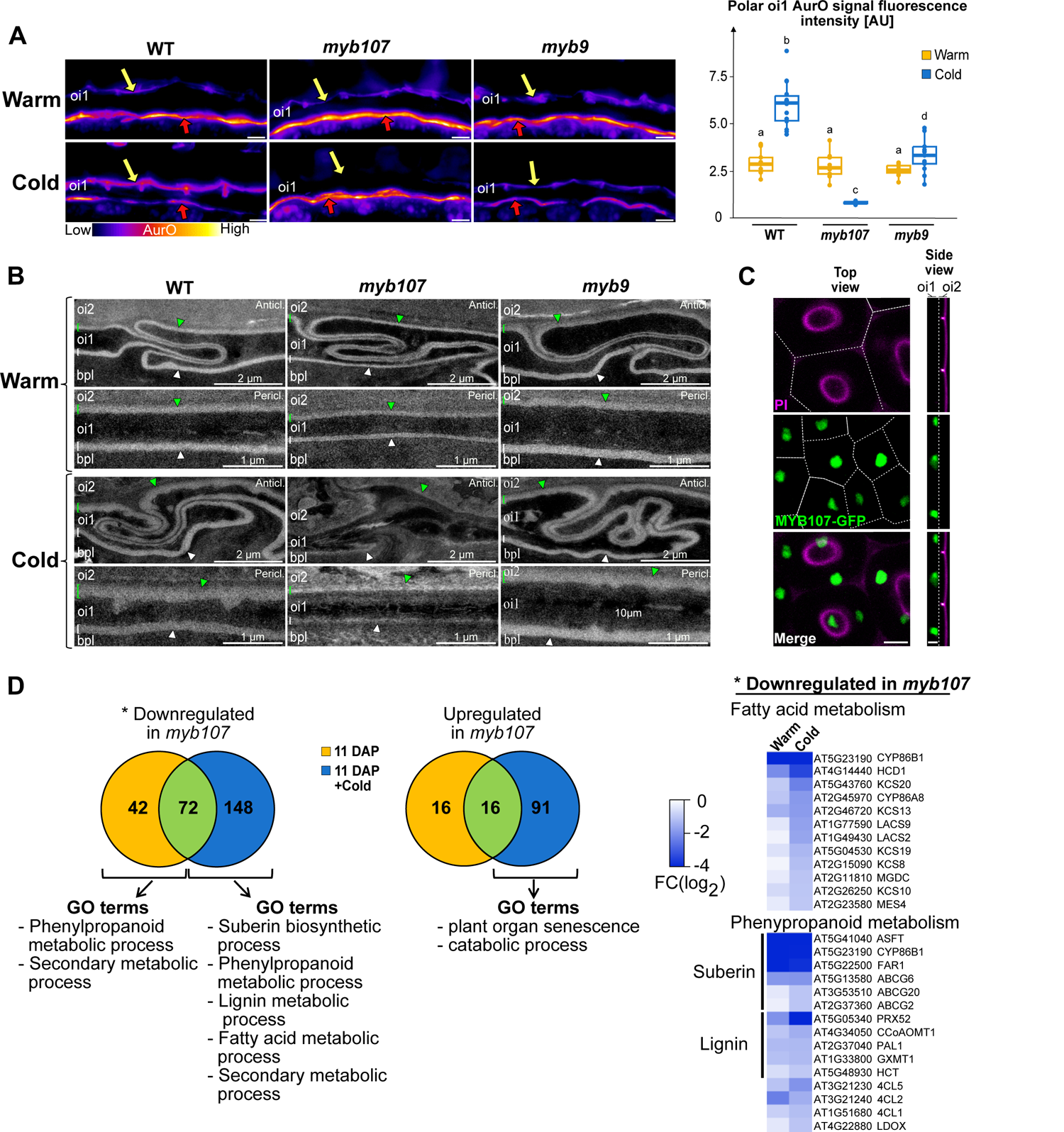
MYB107 is essential for polar oi1 barrier formation in seeds developing under cold temperatures. **A.** Left: AurO staining of WT, *myb107-2* (*myb107*), *myb9-1* (*myb9*) mature Warm and Cold seed sections, as indicated. Yellow and red arrows indicate the AurO signal corresponding to the polar oi1 barrier and the ii1 cuticle, respectively. Bar, 10µm. Right: Box plots of the polar oi1 AurO fluorescence intensity. Statistically significant differences between the different conditions are indicated by different letters as assessed by Kruskal-Wallis or ANOVA test (p < 0.05, n=13-14 seeds per condition). **B.** TEM micrographs showing the linear electron-lucent anticlinal (Anticl.) and periclinal (Pericl.) signal surrounding oi1 cells in WT, *myb107-2* (*myb107*), *myb9-1* (*myb9*) mature Warm and Cold seeds, as indicated. Green and white brackets/arrowheads indicate the outer and inner oi1 electron-lucent signal, respectively. **C.** Confocal images showing propidium iodide (PI) and GFP fluorescence in the seed mature green stage of *myb107*/*pMYB107::MYB107-eGFP* transgenic plants. Bar, 10 µm. **D.** Left: Venn diagrams showing the number of genes that are downregulated or upregulated in *myb107* at 11 DAP and 11 DAP+Cold. The corresponding GO terms are shown below. Right: Heatmap representations of the log2 fold changes (FC) for downregulated genes in *myb107* related to fatty acid and phenylpropanoid metabolism, as indicated.

In *myb9-1* and *myb107-2* Warm seeds, the oi1 electron-lucent signal observed in TEM had a similar appearance relative to that of WT seeds although in *myb107* seeds its width was smaller in both the outer and inner sides (Figs.2B and S2D). In contrast, in *myb9 myb107* Warm seeds, we encountered different categories of electron-lucent linear signals depending on the oi1 cell being examined: the signal was either normal, thinner or almost absent (Fig. S2E). Furthermore, some oi1 cells had an electron-lucent interior. Hence, these results show that *MYB9* and *MYB107* redundantly promote the formation of the oi1 electron-lucent signal, but they also indicate that additional TFs might be involved. In *myb9-*1 Cold seeds, the oi1 electron-lucent signal had a similar appearance and width relative to that of WT seeds even though the AurO signal was weaker (Figs. 2B & S2F). In contrast, the oi1 barrier electron-lucent signal was no longer visible in *myb107-2* Cold seeds. Altogether, these results further support the notion that the AurO fluorescence signal and electron-lucent structure observed in TEM do not correspond to the same oi1 apoplastic components, although the disappearance of the AurO signal and of the electron-lucent signal was always correlated. They also show the importance of MYB107 to form the polar oi1 barrier and the oi1 electron-lucent barrier in Cold seeds.

### *MYB107* expression is enriched in oi1 cells during the mature green stage

We investigated the pattern of *MYB107* expression during seed development. In WT Warm seeds, *MYB107* expression rose from early mature green stage (10 DAP), peaked at 12 DAP, and declined thereafter (Fig. S3A). *myb107* mutant lines transformed with a complementation vector (*myb107/*pMYB107::*MYB107-eGFP*) produced Cold seeds displaying a polar AurO oi1 signal, indicating successful complementation (Fig. S3B). MYB107-eGFP fluorescence was exclusively observed in oi1 cell nuclei during the mature green stage (Fig. 2C). Similarly, a WT plant transformed with a *pMYB9::NLS-3XVenus* transgene also revealed specific expression in oi1 cells during the mature green stage (Fig. S3C). Hence, these results indicate that *MYB107* and *MYB9* are expressed in oi1 cells to promote polar oi1 barrier formation during the mature green stage.

### MYB107 promotes polyester- and lignin-related gene expression under cold temperatures

WT and *myb107* plants cultivated under warm temperatures were transferred to cold (10°C) at 11 DAP, i.e. a time when *MYB107* expression increases to promote polar oi1 barrier formation (Fig. S3A). Total RNA was isolated from dissected seed coats and endosperms at 11 DAP and 48 hours after transfer to cold (11 DAP+Cold) for transcriptome analysis (RNAseq, Table S1). In *myb107* mutants, 114 genes were downregulated at 11 DAP and 220 genes at 11 DAP+Cold, with 72 genes commonly downregulated (Fig. 2D). Additionally, 32 genes were upregulated at 11 DAP and 107 genes at 11 DAP+Cold, with 16 genes commonly upregulated (Fig. 2D). Hence, MYB107 regulates the expression of a larger set of genes upon transfer to cold, which is consistent with its central role to promote polar oi1 barrier formation in Cold seeds.

At 11 DAP, the only genes that could be associated with a gene ontology (GO) category were those downregulated in *myb107* mutants, specifically in the phenylpropanoid and secondary metabolic process categories (Fig. 2D, Table S1). At 11 DAP+Cold, two additional subclasses of phenylpropanoid metabolism emerged: the suberin biosynthetic and lignin metabolism process classes. Additionally, a new category appeared, that of fatty acid metabolism (Fig. 2D, Table S1). Suberin biosynthesis involves fatty acid and phenylpropanoid metabolism whereas lignin biosynthesis involves phenylpropanoid metabolism. More genes associated with fatty acid and phenylpropanoid metabolism were downregulated in *myb107* Cold seeds relative to *myb107* Warm seeds (Fig. 2D, Table S1). These results suggested that the oi1 apoplastic components in Cold seeds contain lignin and suberin. These hypotheses were further investigated below.

### Suberization does not readily account for oi1 cell polarity

TEM sections treated with H_2_O_2_ revealed lamellae in the inner periclinal electron-lucent signal of oi1 cells in WT mature Warm and Cold seeds, consistent with a previous report, but also in the outer periclinal and anticlinal signals, which was not previously reported (Chang-Jun Liu, personal communication) (Fig.3A) (16). Hence, these results support the notion that the electron-lucent signal throughout the oi1 cell contour corresponds to suberization of oi1 cells. We argued above that although the outer electron-lucent signal is rougher and mildly thicker in Cold seeds, the electron-lucent signal does not satisfactorily account for the polar AurO oi1 signal (Fig. 1A & 1B). Hence, polar suberization unlikely explains the polar AurO oi1 signal. Consistent with this notion, no striking differences in the lamellae appearance were observed between Warm and Cold seeds (Fig. 3A). Furthermore, Fluorol Yellow (FY), a fluorescent dye used to stain suberin, revealed an apolar signal surrounding oi1 cells in both WT Warm and Cold seeds (Fig. 3B). The FY signal was moderately stronger in Cold seeds relative to Warm seeds, consistent with the moderate width increase of the electron-lucent signal (Fig. 3B & 1B).

**Figure 3:**
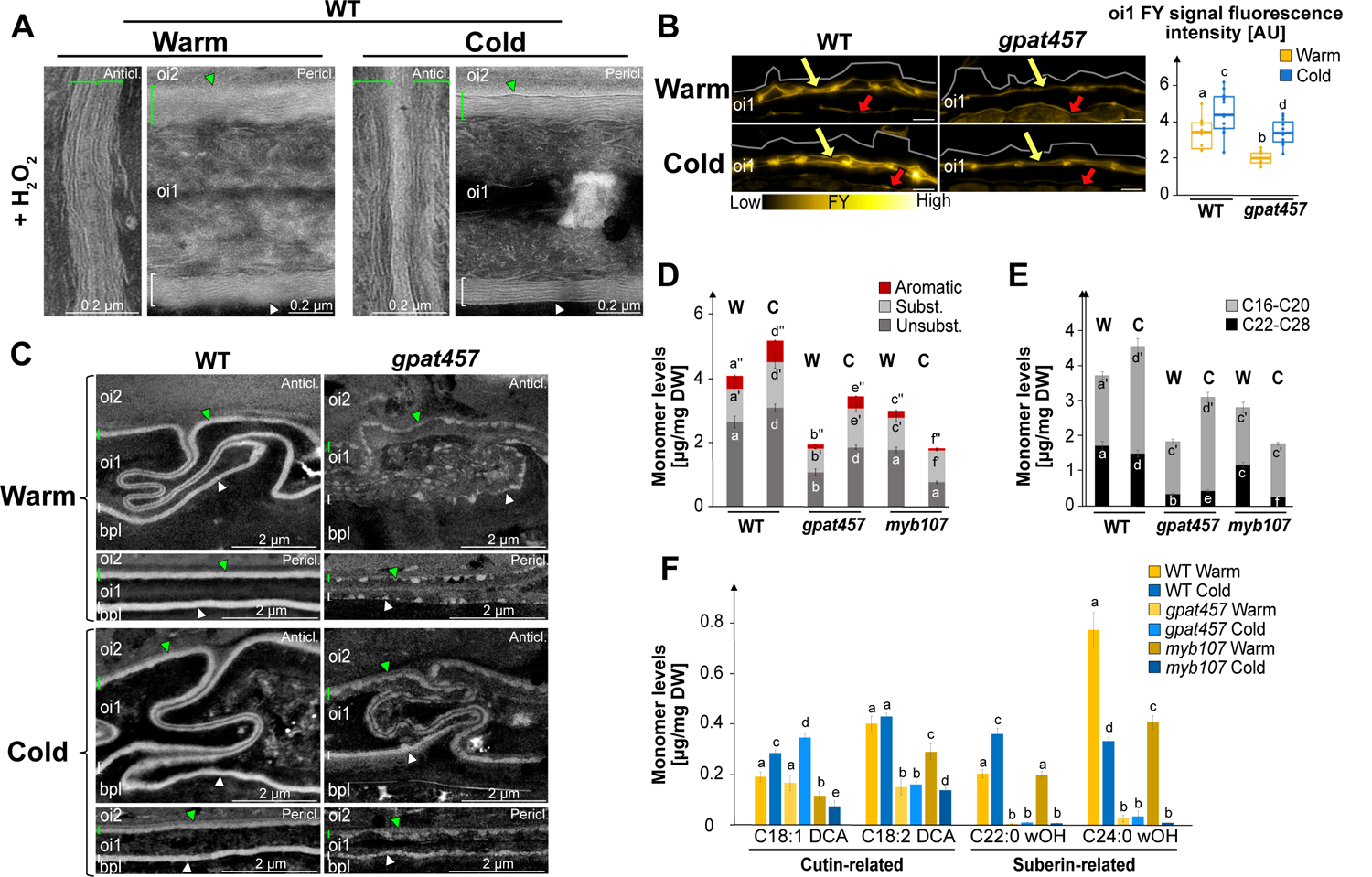
Suberization does not readily account for oi1 cell polarity A. TEM micrographs of WT Warm and Cold seed sections treated with H_2_O_2_ revealing lamellae in the linear electron-lucent anticlinal (Anticl.) and periclinal (Pericl.) signal surrounding oi1 cells. Green and white brackets/arrowheads indicate the outer and inner oi1 electron-lucent signal, respectively. B. Left: Fluorol Yellow (FY) staining of WT and *gpat457* Warm and Cold seed sections, as indicated. Yellow and red arrows indicate the FY signal corresponding to the oi1 barrier and the ii1 cuticle, respectively. Bar, 10µm. Right: Box plots of the oi1 FY signal fluorescence intensity. Statistically significant differences between the different conditions are indicated by different letters as assessed by Kruskal-Wallis or ANOVA test (p < 0.05, n=13-14 seeds per condition). C. TEM micrographs showing the linear electron-lucent anticlinal (Anticl.) and periclinal (Pericl.) signal surrounding oi1 cells in WT and *gpat457* Warm (W) and Cold seeds, as indicated. Green and white brackets/arrowheads as in A. D. Barplots show means, ± SD of aliphatic and aromatic ester-bond monomer levels in WT, *gpat457* and *myb107-2* Warm and Cold mature seeds, as indicated. Subst, substituted; Unsubst., unsubstituted; Aromatic, aromatic. Statistically significant differences between the different genotypes are indicated by different letters as assessed by Kruskal-Wallis or one-way ANOVA test or (p < 0.05, four biological replicates, n = 4). E. Barplots show means, ± SD of aliphatic monomers levels according to their carbon chain length (C16-C20 or C22-C28) in WT, *gpat457* and *myb107-2* Warm (W) and Cold (C) mature seeds, as indicated. Statistics as in D. F. Barplots show means, ± SD of monomer levels associated with cutin or suberin in WT, *gpat457* and *myb107-2* Warm (W) and Cold (C) mature seeds, as indicated. For each monomer, different lowercase letters indicate significant differences, as determined by Krusal-Wallis or one-way ANOVA test, (p < 0.05, n=4).

To further assess suberization in oi1 cells, we studied mutants deficient in polyester biosynthesis. *GPAT5* and *GPAT7* have been associated with suberin production and *GPAT5* was suggested to be expressed in oi1 cells (12, 14, 26). We found that *GPAT5* and *GPAT7* promoter reporter lines (p*GPAT5*::*mCitrine-SYP122, pGPAT7::NLS-GFP*) produced a fluorescent signal specifically in oi1 cells during the mature green stage, indicating that *GPAT5* and *GPAT7* could indeed promote oi1 suberization (Fig. S4A). Nevertheless, *gpat5 gpat7* (*gpat57*) seeds produced a FY signal comparable to that of WT Warm and Cold seeds (Fig. S4B). However, the oi1 electron-lucent signal was severely altered in *gpat57* seeds, appearing as a strand of beads-like electron-lucent depositions (Fig. S4C). Interestingly, in Cold seeds the signal maintained a strand of beads-like appearance mainly on the outer periclinal and outer anticlinal sides of oi1 cells, consistent with a polar apoplastic property of oi1 cells (Fig. S4C). The electron-lucent and FY signals in *gpat57* seeds suggested that other GPATs can promote suberization in oi1 cells. We considered *GPAT4* since a *pGPAT4::NLS-GFP* line also produced a fluorescent signal in oi1 cells during the mature green stage (Fig. S4A). Indeed, the FY signal decreased, although not entirely, in both *gpat4 gpat5 gpat7* (*gpat457*) Warm and Cold seeds (Fig. 3B). However, it remained higher in Cold seeds relative to Warm seeds (Fig. 3B). The oi1 electron-lucent depositions also decreased in *gpat457* compared to WT, maintaining their strand of beads-like aspect, being more abundant in Cold seeds relative to Warm seeds, consistent with the FY results (Fig. 3C). Strikingly, the electron-lucent signal’s polarity was more evident in Cold *gpat457* seeds, with a strand of beads-like appearance on the outer side of oi1 cells and a smoother, linear inner side (Fig. 3C).

Intriguingly, the AurO signal was high in both Warm and Cold *gpat57* seeds, being similar in intensity as WT Cold seeds (Fig. S4B). In contrast, the AurO signal in *gpat457* Warm seeds was similar to WT Warm seeds but its intensity did not increase in Cold *gpat457* seeds (Fig. S4D). Hence, altogether, these results provide further evidence that the electron-lucent and FY signals correspond to polyester depositions in the oi1 barrier, such as suberin; GPAT4, GPAT5 and GPAT7, and likely additional GPATs, are involved in their deposition; oi1 cells have a polar apoplastic property in Cold seeds; and oi1 polyester depositions do not readily explain the polar AurO signal. Cold conditions could induce the polar accumulation of a special type of suberin or of depositions unrelated to suberin that are not detected by FY or TEM but detected by AurO.

### Characterization of cold-induced changes in seed coat polyester content in Arabidopsis

We analyzed esterified lipids in Warm and Cold seeds using GC/MS. Previous studies in *Brassica napus* indicated that the seed coat contains most polyesters (27). De Giorgi et al. found similar results in Arabidopsis, suggesting whole-seed analysis reflects seed coat esterified lipids (28). In WT seeds, polyester monomer content increased modestly in Cold seeds compared to Warm seeds, consistent with TEM and FY results but not with AurO results, further indicating that AurO might detect non-polyester depositions (Fig. 3D).

In *gpat457* Warm seeds and *myb107* Cold seeds, polyester monomer content decreased significantly, correlating with weak or absent oi1 electron-lucent and FY signals (Fig. 3D). This suggests the remaining monomers in these mutants come from other seed coat barriers; or that they are not detected by TEM or FY; or that they are present as diffuse suberin in the oi1 cells interior rather than in the oi1 contour. In *gpat457* seeds, monomers increased in Cold seeds, consistent with TEM and FY results, while in *myb107* seeds, unsubstituted monomer levels remained unchanged, and substituted monomers decreased twofold, suggesting substituted monomers are critical components of the oi1 apoplastic barrier in Cold seeds (Fig. 3D).

In WT Cold seeds, short monomer (C16 to C20) levels increased 1.5-fold, while long monomer (C22 to C28) levels decreased slightly, aligning with TEM and FY results. Similarly, in *gpat457* seeds, short monomers also increased in Cold seeds, consistent with TEM and FY results, whereas long monomers remained low (Fig. 3E). Hence, these results may indicate that cold promotes deposition of short monomers. However, in *myb107* seeds, short monomer levels were unchanged between Warm and Cold seeds, which invalidates our first hypothesis or suggests that cold promotes short monomer accumulation elsewhere in the seed coat (Fig. 3E). Long monomer levels were significantly lower in *gpat457* seeds compared to WT, and in *myb107* Cold seeds, long monomer levels were the lowest measured, correlating low long monomer levels with a low oi1 AurO signal (Fig. 1E). Hence, although an increase in the oi1 barrier AurO signal in WT Cold seeds relative to WT Warm seeds could not be correlated with an increase in long monomer levels, these findings suggest long monomers are components of the cold-induced polar oi1 barrier and that AurO detects other compounds besides polyesters whose deposition is promoted by cold in oi1 cells. Therefore, it is unclear what is the relative contribution of short and long monomers in the oi1 polyester depositions.

Individual monomer analysis showed that C22:0-OH and C24:0 wOH levels, associated with suberin, were reduced in *myb107* and *gpat457* Cold seeds, while C18:1 DCA and C18:2 DCA levels, associated with cutin, were less affected (Fig. 3F, S4E & S4F) (29). This suggests suberin is a major component of the oi1 apoplastic barrier in Cold seeds. Suberin can be associated with fatty acids bearing aromatic groups derived from the phenylpropanoid pathway such as coumarate, ferulate, or sinapic acid (30). Ferulate cis and ferulate trans levels increased in WT Cold seeds but were significantly lower in *gpat457* and *myb107* seeds, suggesting AurO might detect compounds containing aromatic acids like ferulate (Fig. S4F). The measurements with *myb107* seeds are consistent with transcriptomic data showing that MYB107 regulates the expression of phenylpropanoid metabolism genes in Warm and particularly Cold seeds (Fig. 2D).

### Cold promotes the polar deposition of lignin or lignin-like polymers in oi1 cells

Previous studies suggested lignin presence in the seed abscission zone, but whether lignin is present elsewhere in the seed coat is unclear (20). Using the Wiesner test, a purple signal appeared between the oi1 and oi2 cell layers in WT Cold seeds but was less visible in Warm seeds (Fig. S5A). Furthermore, Basic Fuchsin (BF), a stain that binds to lignin, revealed a weak polar signal in oi1 cells of WT Warm seeds, which increased markedly in Cold seeds, arising from the outer periclinal and anticlinal parts of oi1 cells, similar to the polar AurO signal (Fig.4A). Hence, these results indicate that cold may promote the polar deposition of lignin or lignin-like polymers in oi1 cells.

**Figure 4:**
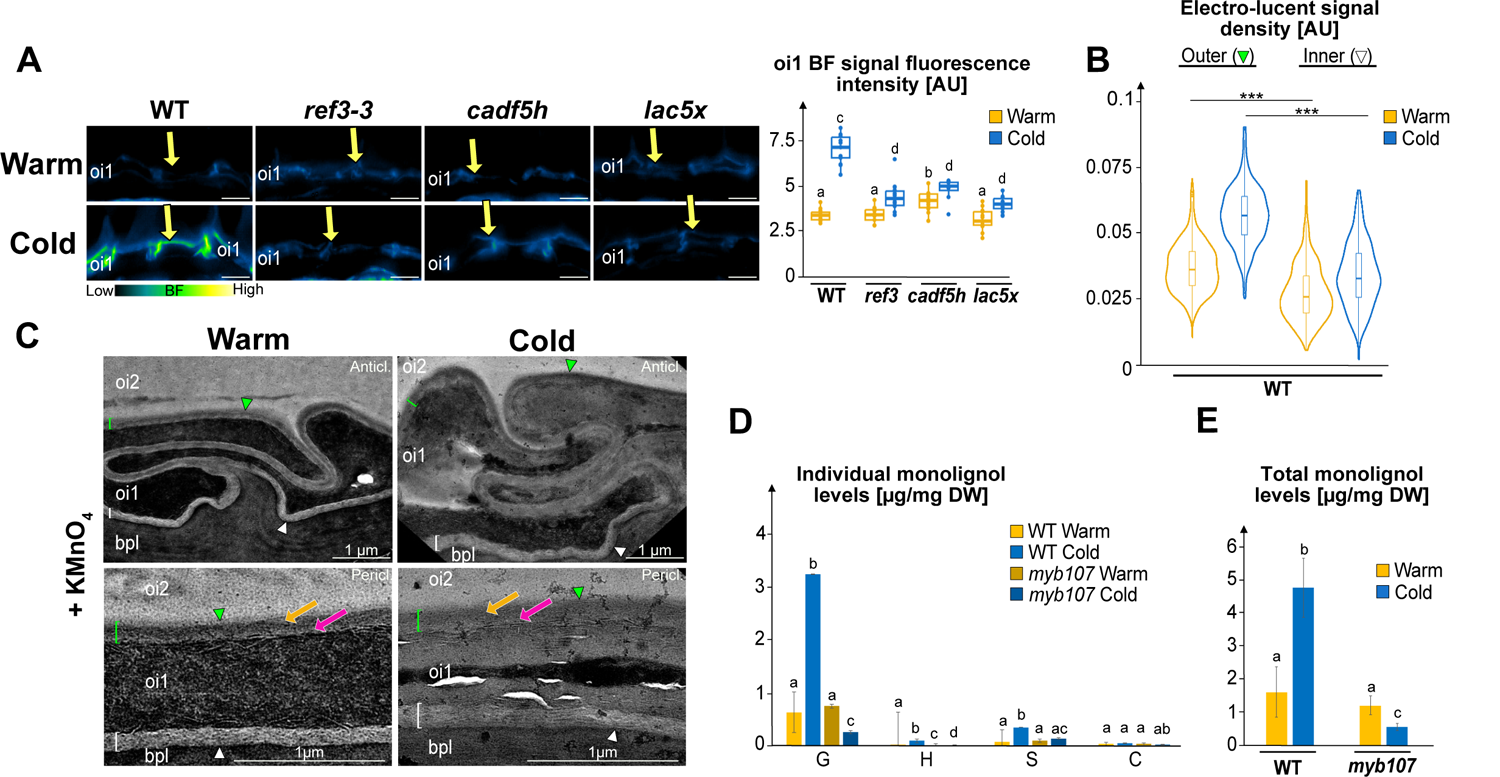
Cold promotes the polar deposition of lignin or lignin-like polymers in oi1 cells A. Left: Basic Fuchsin (BF) staining of WT, *ref3-3*, *cadf5h* and *lac5x* mature Warm and Cold seed sections. Yellow arrow indicates the BF signal corresponding to the polar oi1 barrier. Bar, 10µm. Right: Box plots of the oi1 BF signal fluorescence intensity. Statistically significant differences between the different conditions are indicated by different letters as assessed by a Kruskal-Wallis or ANOVA test (p < 0.05, n= 8-14 seeds per condition). B. Violin plots of the outer and inner electro-lucent signal electron density in WT Warm and Cold TEM sections treated with KMnO_4_. Statistical analysis as assessed by Kruskal-Wallis test (***p< 0.001). C. TEM micrographs of WT Warm and Cold seed sections treated with KMnO_4_ revealing revealing higher electron density in the outer anticlinal (Anticl.) and periclinal (Pericl.) electron lucent signal surrounding oi1 cells. Green and white brackets/arrowheads indicate the outer and inner oi1 linear signal, respectively. Orange and pink arrows show regions of higher and lower electron density in the outer oi1 linear signal. D. Abundance of diagnostic thioacidolysis monomers specifically released from -O-4-linked p-hydroxyphenyl (H), guaiacyl (G), syringyl (S), and catechyl (C) lignin units in WT and *myb107* Warm and Cold seeds. Values represent means ± SD, n = 4. For each monomer, different lowercase letters indicate significant differences, as determined by Krusal-Wallis or one-way ANOVA test (p < 0.05, four biological replicates, n=4). E. Total lignin monomer abundance released by analytical thioacidolysis in WT and *myb107* Warm and Cold seeds. Statistics as in D.

We further assessed this claim genetically. CINNAMYL ALCOHOL DEHYDROGENASEs (CADs) reduce cinnamaldehydes into cinnamyl alcohols in monolignol biosynthesis, while FERULATE-5-HYDROXYLASEs (F5H) convert guaiacyl monolignol to syringyl monolignol. The quadruple mutant *cadf5h*, lacking *CAD4*, *CAD5*, *F5H1*, and *F5H2*, is defective in monolignol biosynthesis and showed decreased BF signals in Cold seeds compared to WT Cold seeds (Fig.4A) (31). We also analyzed the *reduced epidermal fluorescence 3* (*ref3-3*) mutant, a hypomorphic allele of *CINNAMATE-4-HYDROXYLASE* (*C4H*), and the quintuple *laccase 1,3,5,13,16* (*lac5x*) mutant, defective in monolignol polymerization (31–33). Both *ref3-3* and *lac5x* also displayed reduced BF signals in Cold seeds (Fig.4A). These results provide genetic support to the notion that cold promotes the accumulation of lignin or lignin-like depositions in oi1 cells.

We acquired TEM images of WT mature seed sections treated with potassium permanganate (KMnO_4_), which enables visualizing lignin depositions as electron-dense depositions (34–36). This showed increased electron density in the outer periclinal and outer anticlinal parts of the oi1 barrier while the inner parts remained electron-lucent (Fig. 4B-C). Cold further increased the electron density in the outer oi1 barrier relative to the inner oi1 (Fig. 4B-C). In both Warm and Cold WT seeds, the higher electron density was observed in the outer part of the outer oi1 barrier whereas its inner part retained an electron-lucent character like that of the inner oi1 signal (Fig. 4C). Altogether, these data further support the notion that oi1 cell walls are polarly lignified while suberized throughout their contour. They also support the notion that cold enhances polar lignification of oi1 cells.

Transgenic lines with NLS-GFP reporters driven by *PHENYLALANINE AMMONIA-LYASE 1* (*PAL1*), *PAL2*, and *C4H/REF3* promoters produced fluorescent signals in oi1 cells at the mature green stage, while a *PAL4* promoter did not display a signal (Fig. S4A). These data suggest that oi1 cells autonomously control their lignin depositions.

To chemically verify lignin depositions in the Arabidopsis seed coat we performed analytical thioacidolysis, which selectively cleaves alkyl aryl ether bonds, releasing the diagnostic β-O-4-linked p-hydroxyphenyl (H), guaiacyl (G), and syringyl (S) lignin units/ monomers. The seed lignin is predominantly composed of G units, with lower amounts of S-type monomers (Fig. 4D). H units and coniferyladehyde were also detected but are near the detection limit and partially absent in *myb107* (Fig. 4D). Interestingly, caffeyl alcohol derived catechyl units (C), known from other seed coat lignins were also detected in minor amounts (Fig. 4D) (37).

When quantifying monolignols in WT and *myb107* seeds using GC-MS we determined similar total monolignol levels in WT and *myb107* Warm seeds (Fig. 4E). Strikingly, total monolignols levels increased by threefold in WT Cold seeds relative to Warm seeds, whereas twofold reduced levels were detected in *myb107* Cold seeds relative to Warm seeds. The cold-induced lignin deposition is mainly a result from increases in G, S and H monomers. Furthermore, consistent with the AurO results and monolignol measurements, a markedly reduced BF signal was found in *myb107* Cold seeds (Fig. S5C). These results support the notion that cold stimulates through MYB107 the deposition of a lignin polymer in the seed coat resembling that found in other parts of the plant.

### Defects in the oi1 barrier correlate with defects in seed coat impermeability and seed dormancy

We investigated whether defects in the oi1 barrier are associated with defects in seed permeability and dormancy. Assessing the permeability of individual seed coat cell integumental layers is challenging. Previous reports showed that *myb107* and *myb9* mature Warm seeds have higher permeability to Tetrazolium Red, consistent with a defective seed coat in these mutants. Tetrazolium Red permeability assesses the entire seed coat, including the ii1 cuticle tightly associated with the endosperm, which makes it uncertain whether the higher permeability reflects defects in the seed coat’s outer layers (8). We therefore sought to develop a protocol more specific to the outer layers of the seed coat. The outer integuments are external to the bpl and ii1 seed coat layers, which contain oxidized tannins giving the seed coat its brown color. Treating seeds with sodium hypochlorite leads to tannin breakdown causing the seed coat to lose color, appearing as white spots (partially bleached) or white (fully bleached) seeds. Hence, whitening can be used as a readout to assess whether defects in the outer integuments correlate with higher seed coat permeability to sodium hypochlorite. *myb107* Warm seeds, with a thinner electron-lucent oi1 barrier, had 48% spotted seeds and 10% white seeds, while WT Warm seeds had no spotted seeds and 7% white seeds (Fig. 5A). *myb9* Warm seeds, with no oi1 barrier defects, showed no color defects. *myb107 myb9* Warm seeds, which are highly defective in oi1 apoplastic depositions, had 76% white seeds and 18% spotted seeds. Similar results were obtained with *myb107* Cold seeds (Fig. 5B), whereas WT and *myb9* Cold seeds had no color defects.

**Figure 5:**
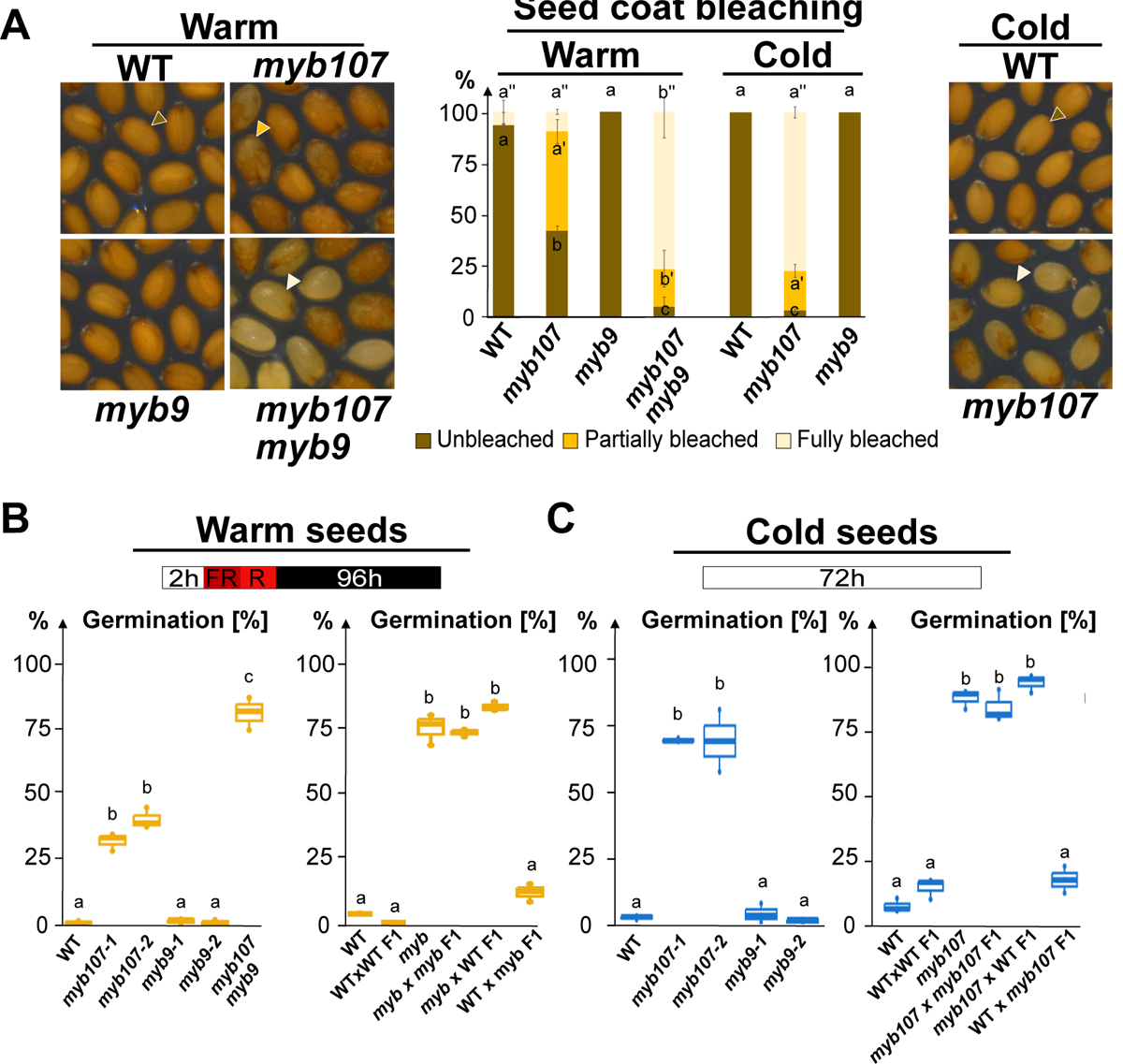
Defects in the oi1 barrier correlate with defects in seed coat impermeability and seed dormancy. **A.** Left: Pictures showing WT, *myb107-2* (*myb107*)*, myb9-1* (*myb9*) and *myb107-2 myb9-1* (*myb107 myb9*) mature Warm seeds after 15 min incubation with sodium hypochlorite. Right: Pictures showing WT and *myb107-2* (*myb107*) mature Cold seeds after 15 min incubation with sodium hypochlorite. Brown, orange and white arrowheads show unbleached, partially bleached and fully bleached seeds, respectively. Center: Histograms showing the percentage of each category of bleaching in Warm and Cold mature seeds of various genotypes, as indicated. Statistically significant differences between the different genotypes are indicated by different letters as assessed by a two-way ANOVA test followed by a post-hoc Tukey test, (p < 0.05, n= 60-100 seeds per condition). **B.** Schematic showing the suboptimal germination protocol consisting of imbibing seeds (15 days after harvesting) for 2h in white light, followed by a 5min far red (FR) pulse immediately followed by a 5 min red (R) pulse and 96h hours dark incubation before assessing germination. Left: Box plots showing germination percentages of Warm mature seeds of various genotypes, as indicated. Right: Reciprocal crosses showing that the low dormancy of *myb107-2 myb9-1* (*myb*) mature Warm seeds is maternally inherited. Germination was assessed. Statistics as in A. **C.** Shematic showing the optimal germination protocol consisting of imbibing seeds for 72h in white light before assessing germination. Left: Box plots showing germination percentages of Cold mature seeds (36 days after harvesting) of various genotypes, as indicated. Right: Reciprocal crosses showing that the low dormancy of *myb107-2* (*myb107*) mature Cold seeds (67 days after harvesting) is maternally inherited. Statistics as in A.

We assessed whether oi1 barrier defects are linked to seed dormancy defects. WT Col-0 accession Warm seeds have low dormancy when assessed under optimal germination conditions, which complicates the assessment of dormancy defects in mutants. Using suboptimal germination conditions, which enhance germination arrest at low dormancy levels, only a few percent of WT Warm seeds germinated after 5 days with a far red (FR) pulse followed by a red (R) pulse and incubation in darkness (Fig. 5B) (24). Similar results were obtained with *myb9-1* and *myb9-2* Warm seeds (Fig. 5B). In contrast, over 30% germination occurred with *myb107-1* and *myb107-2* Warm seeds, and 81% with *myb107 myb9* Warm seeds (Fig. 5B). F1 Warm seeds from *myb107 myb9* plants pollinated with WT pollen showed 83% germination whereas only a few percent germination was observed in the reciprocal cross, indicating maternally inherited low dormancy (Fig. 5B).

WT Col-0 accession Cold seeds, which have increased seed dormancy levels, and mutant Cold seeds were tested under standard conditions with white light. WT and *myb9* Cold seeds did not germinate when tested 36 days after harvest, while 60-75% of *myb107* mutants did (Fig. 5C). Low dormancy was again maternally inherited (Fig. 5C).

Altogether, these results establish a correlation between oi1 barrier defects and low seed coat permeability and low seed dormancy. They are also consistent with the notion that cold promotes dormancy by promoting polar oi1 lignification and suberization throughout the entire oi1 cell boundary.

## Discussion

Here, we presented genetic, histological, and biochemical analyses suggesting that cold induces polar lignification in the oi1 cell layer of the Arabidopsis seed coat. The S/G ratio in WT seed lignin (0.11-0.12) is more similar to root (0.21) and Casparian strip lignin (0.09) than to stem lignin (0.67) (31, 38, 39). Nevertheless, the H/G/S monomeric composition resembles an angiosperm lignin polymer that is similar or closely related to classical lignin found in other cell types in the plant (40). Although lignin presence in Arabidopsis seed coat was not well-documented, previous reports indicate its presence in Brassicaceae (41, 42). Prior work hints at lignin production in Arabidopsis oi1 cells, with TT10, a laccase-encoding gene, being expressed in these cells, and *tt10* mutants showing reduced lignin content (21, 22). To our best knowledge, this is the first report showing polar lignification in Arabidopsis seed coat in response to cold during seed development. Lignin in the Arabidopsis seed coat might have been overlooked as it is significantly deposited only in cold conditions. Previous studies reported the presence of lignin in the seed coats of Orchids and Cacteaceae (37, 43). It remains to be investigated whether cold promotes seed coat lignification in other species, including Brassicaceae.

The oi1 layer was previously proposed to produce suberin, however suberin presence was only indirectly ascertained by the observation of lamellae on the inner side within the electron-lucent linear signal (13, 16). Here, we presented independent lines of evidence further supporting this proposition: 1) presence of lamellae throughout the oi1 cell surface, 2) an apolar FY signal around oi1 cells and 3) an altered electron-lucent signal in *gpat57* and *gpat457* oi1 cells. Interestingly, we found that GPAT4, linked to cutin biosynthesis, also contributes to oi1 barrier formation. Recent work showed that *GPAT4*, redundantly with *GPAT6* and *GPAT8*, that are also associated with cutin biosynthesis, is required for root suberization (44). Our results suggest that additional *GPATs* promote oi1 suberization as electron-lucent material increased in Cold *gpat457* seeds. The altered, “strand of beads-like”, electron-lucent depositions in *gpat* mutants described here resemble the reported alterations in suberin deposition, characterized by irregular amorphous structures and gaps in suberin coverage (44).

We confirmed that MYB107 promotes the expression of polyester, lignin, and phenylpropanoid genes, especially in cold seeds where the number of lignin and phenylpropanoid biosynthesis genes regulated by MYB107 increases. We also showed that MYB107 plays a predominant role to promote both suberization and lignification of oi1 cells in Cold seeds, whereas both MYB107 and MYB9 redundantly promote these processes in Warm seeds likely with additional factors. Transgenic reporter lines indicated that *MYB107* and genes encoding enzymes of the phenylpropanoid pathway are specifically expressed in oi1 cells, indicating that oi1 cells autonomously promote their lignification in response to cold.

Upon transfer to cold at 12 DAP, *MYB107* expression further increased after 2 days, suggesting that cold promotes MYB107 accumulation (Fig. S3A). Hence, cold could enhance polar lignin deposition by increasing *MYB107* expression. Furthermore, cold also slows seed development, likely prolonging the seed coat’s life span, which could provide more time for oi1 lignification and suberization. The mechanism of polar lignin deposition in oi1 cells remains to be investigated. Similar mechanisms to Casparian strip formation in the root endodermis might be involved, involving CASP proteins that localize lignin biosynthesis enzymes such as peroxidases and laccases. The *CASP-LIKE* gene *CASPL1D2* (AT3G06390) was downregulated in *myb107* Cold seeds (11DAP + Cold) whereas *CASPL1C1* (AT4G03540) is downregulated in both *myb107* Warm and Cold seeds (11DAP & 11DAP+Cold), suggesting that they could play a role in oi1 polar lignin deposition (Table S1).

Spatial control of lignification in the context of the Casparian strip formation may be necessary to prevent ROS reactions from spreading within endodermal cells and compromise their viability. Since oi1 cells are destined to die, lignification in these cells may involve mechanisms similar to xylem lignification, which is linked to cell death. Polarization may involve polar deposition of lignin monomers, monolignol transporters, dirigent proteins or tyrosine-rich cell wall proteins associated with lignin deposition (45–50). Hydroxycinnamate esters in certain cell wall polysaccharide domains could also play a role (51). A combination of these mechanisms is possible. The “pointed hat” pattern of the electron-lucent anticlinal signals indicates oi1 cell polarization (Fig. S1D and Fig. 6). The boundary of oi1 lignification is located at the tip of the “pointed hat”, i.e. where anticlinal adjacent oi1 cell walls separate inwardly after running in close proximity with each other more outwardly (Fig. 6). Close proximity may serve as a scaffold for initial monolignol deposition, also ensuring a tight seal between cells before spreading outward towards the outer periclinal cell wall (Fig. 6). The polarization in *gpat457* Cold seeds, with a strand of beads-like outer signal and smoother inner signal, suggests coordinated lignification and suberization of the outer oi1 barrier (Fig. 3C).

**Figure 6:**
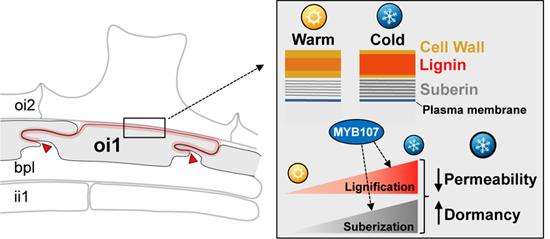
Model for the temperature-dependent polar lignification of a seed coat suberin layer promoting dormancy in Arabidopsis thaliana In response to cold during seed development MYB107 promotes the polar lignification of oi1 cells, i.e. a region shown in red starting at the tip of a prototypical “pointed hat” structure, indicated by red arrow heads, where the cell walls of two adjacent oi1 anticlinal cell walls come in close proximity and form outwardly convolutions before separating (see also Fig. S1D). MYB107 also promotes suberization throughout the oi1 cell contour in response to cold. Increased oi1 cell layer lignification and suberization leads to lower seed coat permeability thus delaying the release of seed dormancy.

We found that defects in the oi1 barrier correlate with high seed coat permeability and low dormancy, indicating its role to promote these seed properties. We speculate that increased suberization and lignification, induced by cold, enhance dormancy by making the seed coat more impermeable thus delaying oxidation events that release dormancy (Fig. 6). Cold temperatures during seed development increase dormancy in many species. Pre-harvest sprouting, the premature germination of seeds before harvest, reduces seed quality and causes economic losses in crops like wheat, barley, and rice. Investigating whether cold promotes seed coat lignification and involves MYB homologs in other species is not only interesting from a biological perspective but it could also lead to developing crops with increased seed coat hardiness and reduced PHS.

## Material and methods

### Plant material and growth conditions

All genotypes used in this study were in the Col-0 background. *myb107-1* (SAIL_242_B04), *myb107-2* (SALK_203615), *myb9-1* (SALK_149765C) and *myb9-*2 (CS463394) were previously described (16, 17). The double mutant *myb107myb9* was generated by crossing *myb107-2* with *myb9-1. gpat457* was generated by crossing *gpat4-1* (SALK_106893), *gpat5-1* (SALK_018117) and *gpat7-3* (SALK_064514), which were previously described (12, 26, 52). *cad4cad5f5h-1f5h-2, lac5x (lac1,3,5,12,16)* and *ref3-3* were as previously described (31–33). The *pGPAT5*::*mCitrine-SYP122*, *pC4H::3xNLS-Venus, pPAL1::3xNLS-Venus, pPAL2::3xNLS-Venus and pPAL4::3xNLS-Venus* lines were as previously described (53, 54). Oligonucleotides used for genotyping are listed in Table S2.

All plants were grown under standard conditions as described (28). Cold was applied upon bolting by transferring plants to a cold chamber (10-13°C, 16 h/8 h day/night photoperiod, light intensity of 120-130 µmol/m^2^s, humidity of 60-65%). Seed batches were harvested on the same day from plants grown side by side under identical environmental conditions. A full description of materials and methods is included in the Supplementary Information (SI) Appendix.

## Acknowledgements

We thank the following colleagues for kindly sharing seed material: Niko Geldner (*cad4cad5f5h-1f5h-2*, *lac5x* (*lac1,3,5,12,16*), *pC4H::3xNLS-Venus*, *pPAL1::3xNLS-Venus*, *pPAL2::3xNLS-Venus* and *pPAL4::3xNLS-Venus*) and Roman Ulm (*ref3-3*). Fabienne Cléard is thanked for helping with cloning and for generating the modified pFR7m34GW for Gibson assembly. We thank Mylene Docquier and members of the Genomics Platform of the Institute of Genetics and Genomics (iGE3) at the University of Geneva for help with RNAseq experiments. We thank the Electron Microscopy Facility of University of Lausanne for help with KMnO_4_-treated seed sections. This work was supported by grants from the Swiss National Science Foundation (No 310030_219559) and by the State of Geneva.

