## Supplementary Figures for "Temperature-dependent polar lignification of a seed coat suberin layer promoting dormancy in Arabidopsis thaliana"

#### Supplementary Figure 1

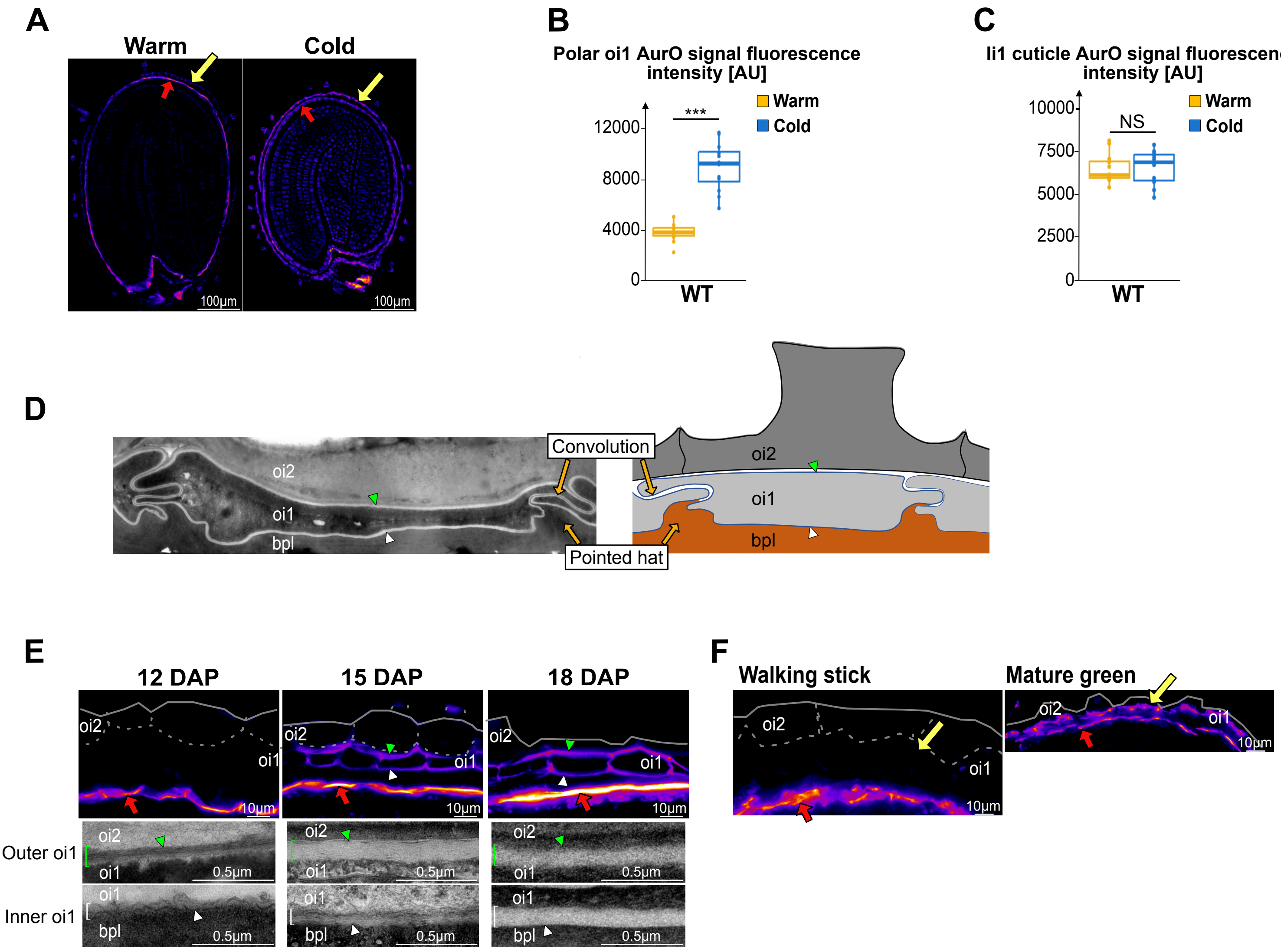


**Supp. Figure 1: Identification of a polar oi1 barrier reinforced by cold during seed development**

**A**. AurO staining of WT Warm and Cold seed sections. Yellow and red arrows indicate the AurO polar oi1 signal and the ii1 cuticule, respectively.
**B**. Box plots of the polar oi1 AurO signal fluorescence intensity in arbitrary units [AU]. Statistical differences assessed by Wilcoxon test (***p< 0.001, n=12 seeds per condition).
**C**. Box plots of the ii1 cuticle Auro signal fluorescence intensity. Statistical differences assessed by ANOVA (NS: not significant p>0.05, n=12 seeds per condition).
**D.** Schematic describing the prototypical convolution and "pointed hat" structure formed by the oi1 linear electro-lucent signal. TEM picture is taken from Fig.1B.
**E.** AurO staining and TEM micrographs showing the AurO fluorescent oi1 signal and the linear signal surrounding oi1 cells in sections of developing Warm seed at 12, 15 and 18 days after pollination (DAP), as indicated. Green arrowhead, outer oi1 barrier; white arrowhead, inner oi1 barrier, red arrow indicates ii1 cuticle, green and white brackets indicate outer and inner oi1 TEM linear signal, respectively.
**F.** AurO staining showing the AurO fluorescent oi1 signal in sections of developing Cold seeds (13˚C) in the walking stick and mature green stages, as indicated. Yellow arrow indicates the AurO signal from the oi1 barrier, red arrow indicates ii1 cuticle.

#### Supplementary Figure 2

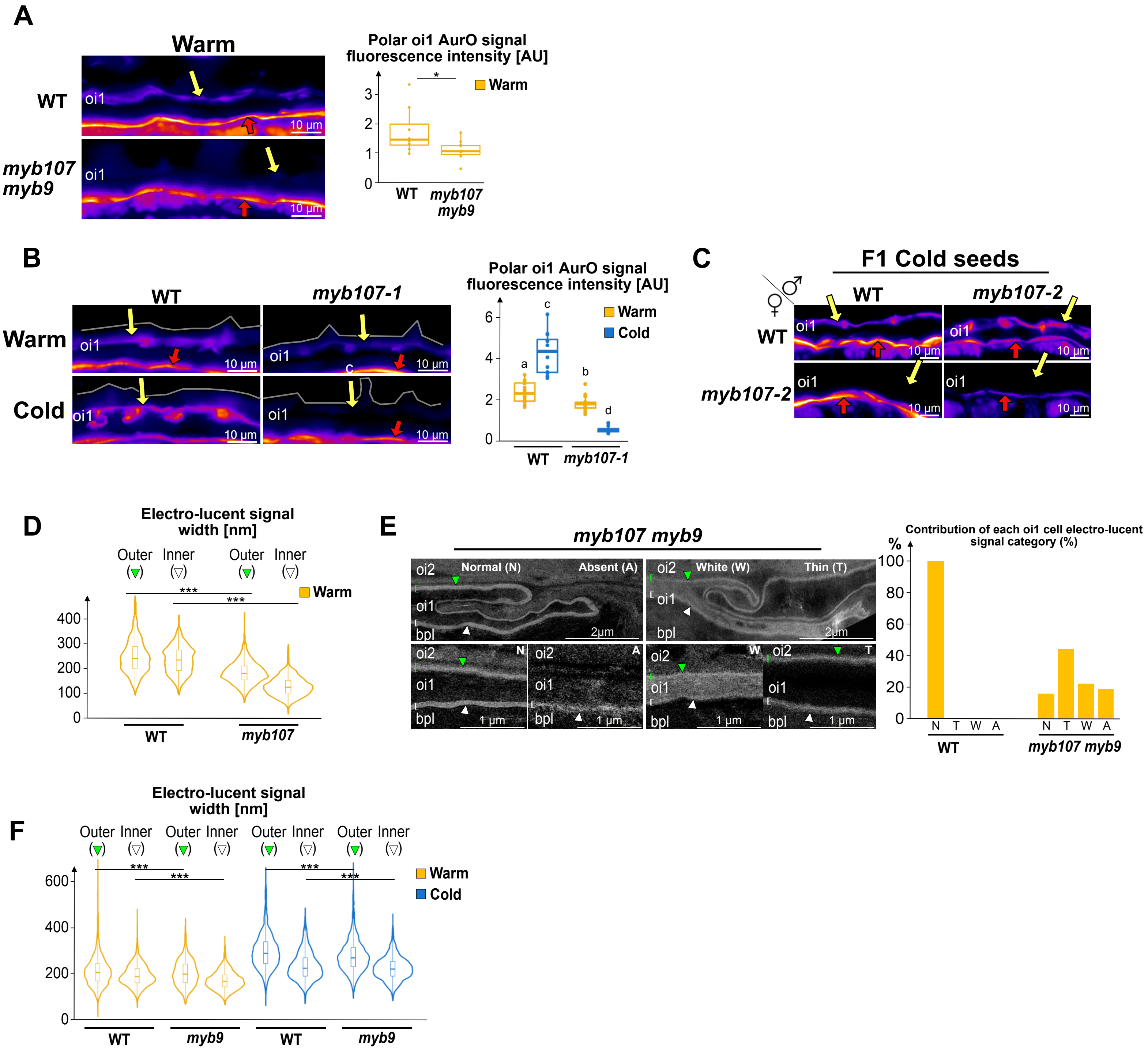


**Supplementary figure 2: MYB107 is essential for polar oi1 barrier formation in seeds developing under cold temperatures**

**A**. Left: AurO staining of WT and *myb107-2 myb9-1* (*myb107 myb9*) mature Warm seed sections. Right: Box plots of the polar oi1 AurO signal fluorescence intensity in arbitrary units [AU]. Statistical differences assessed by ANOVA test (*p< 0.05, n=12 seeds per condition). Yellow and red arrows indicate the AurO signal corresponding to the polar oi1 barrier and the ii1 cuticle, respectively.

**B**. Left: AurO staining of WT and *myb107-1* mature Warm seed sections, as indicated. Right: Box plots of the polar oi1 AurO signal fluorescence intensity in arbitrary units [AU]. Statistical differences assessed by Kruskal-Wallis or ANOVA test (p< 0.05, n=12 seeds per condition). Yellow and red arrows indicate the AurO signal corresponding to the polar oi1 barrier and the ii1 cuticle, respectively.

**C**. AurO staining of F1 Cold seeds arising from crosses between WT and *myb107* plants, as indicated. Yellow and red arrows indicate the AurO signal corresponding to the polar oi1 barrier and the ii1 cuticle, respectively.

**D**. Violin plots of the outer and inner electro-lucent signal width in WT and *myb107-2* Warm seeds shown in Fig. 2B. n=6 cells (3 seeds, 2 cells per seed), statistical analysis as assessed by Kruskal-Wallis test (***p< 0.001).

**E**. Left. TEM micrographs of *myb107 myb9* Warm seeds showing the different categories of oi1 electron-lucent linear signal: Normal (N), Absent (A), White (W) and Thin (T). Right. Histrograms show the percentage contribution of each oi1 cell electron-lucent signal category. For each genotype, 64 oi1 cells distributed over 3 seeds were assessed.

**F**.Violin plots of the outer and inner electro-lucent signal width in WT and *myb9* Warm and Cold seeds shown in Fig. 2B. n=6 cells (3 seeds, 2 cells per seed), statistical analysis as assessed by Kruskal-Wallis test (***p< 0.001).

### Supplementary Figure 3

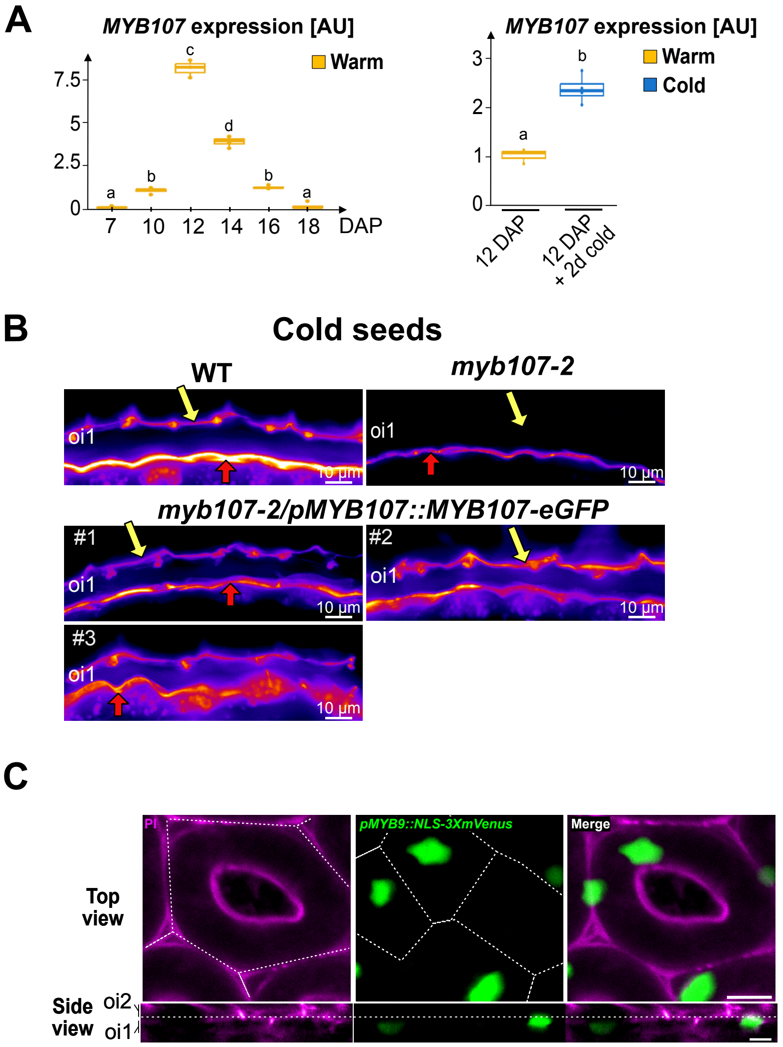


**Supp. Figure 3: MYB107 is essential for polar oi1 barrier formation in seeds developing under cold temperatures**

**A**. Left: Box plots show the relative *MYB107* mRNA accumulation WT Warm seed development at different days after pollination (DAP), as indicated. Right: Relative *MYB107* mRNA accumulation in WT Warm seeds at 12 DAP and after a two days upon transfer to cold. Expression levels were normalized to those of CDK2. n=3-4 technical replicates. Statistically significant differences between the different conditions are indicated by different letters as assessed by a Kruskal-Wallis or ANOVA test ($p < 0.05$).

**B**. AurO staining of WT, *myb107-2* (*myb107*) and *myb107-2*/*pMYB107::MYB107-eGFP* Cold seed sections, as indicated. Yellow and red arrows indicate the AurO signal corresponding to the polar oi1 barrier and the ii1 cuticle, respectively. Three independent complementation lines are shown (#1, #2 and #3).

**C.** Confocal images showing propidium iodide (PI) and GFP fluorescence in the seed mature green stage of *pMYB9:NLS-3XVenus* transgenic plants. Bar, 10 µm.

### Supplementary Figure 4

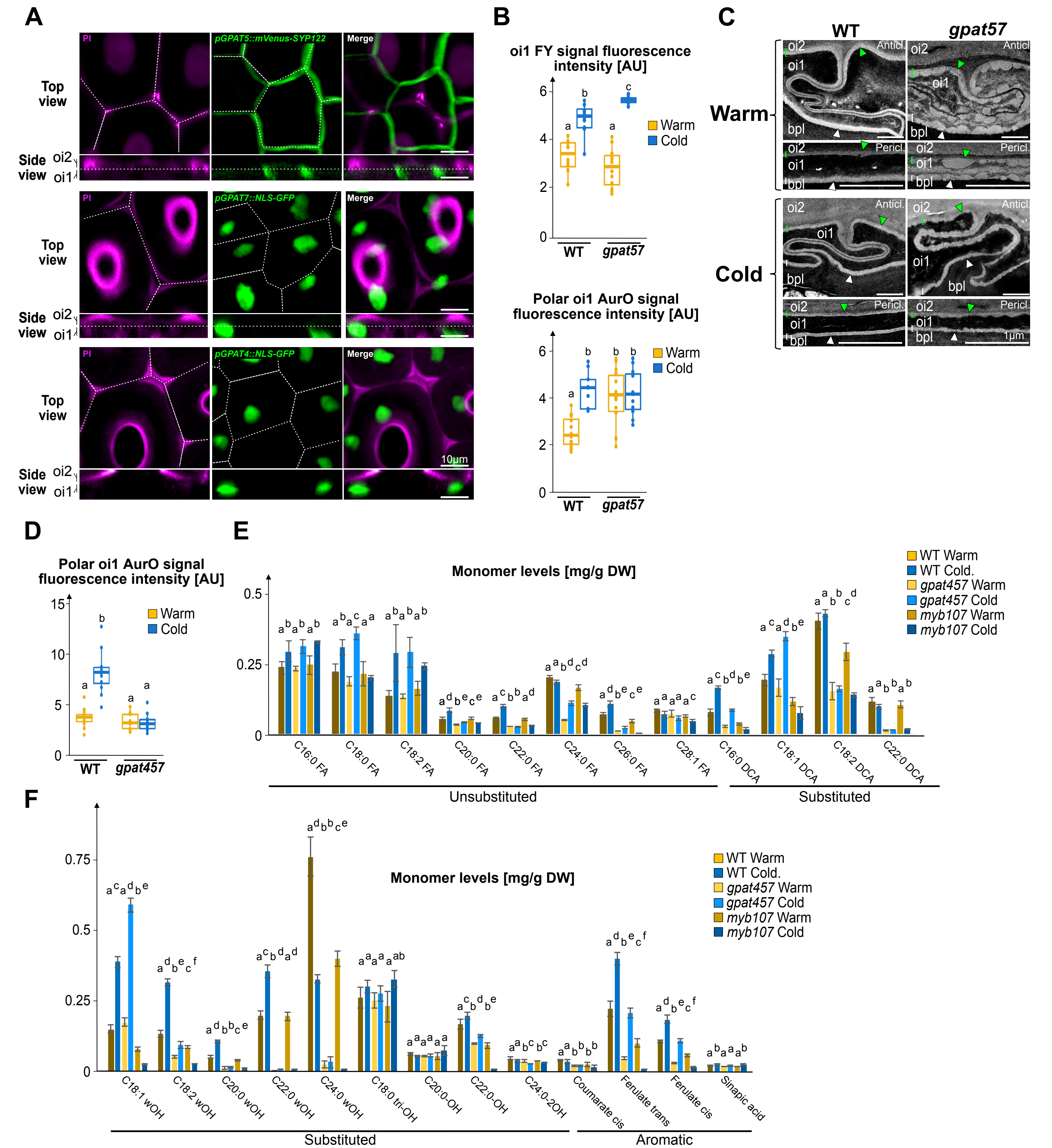


**Supplementary Figure 4: Suberisation does not readily account for oi1 cell polarity**

**A.** Confocal images showing propidium iodide (PI) and GFP or Venus fluorescence in the seed mature green stage of *pGPAT5::mVenus-SYP12*, *pGPAT7::NLS-GFP* and *pGPAT4::NLS-GFP* transgenic plants, as indicated. Bar, 10 µm.

**B.** Top: Box plots of the oi1 FY signal fluorescence intensity in Fig. 3B. Statistically significant differences between the different conditions are indicated by different letters as assessed by a Kruskal-Wallis or ANOVA test ($p < 0.05$, n=12-13 seeds per condition). Botton: Box plots of the polar oi1 AurO signal fluorescence intensity in Fig. 3B. Statistically significant differences between the different conditions are indicated by different letters as assessed by an ANOVA test ($p < 0.05$, n=12-14 seeds per condition).

**C.** TEM micrographs showing the linear electron-lucent anticlinal (Anticl.) and periclinal (Pericl.) signal surrounding oi1 cells in WT and *gpa57* Warm (W) and Cold seeds, as indicated. Green and white brackets/arrowheads indicate the outer and inner oi1 electro-lucent signal, respectively.

**D.** Box plots of the polar oi1 AurO signal fluorescence intensity in WT and *gpat457* Warm and Cold seeds, as indicated. Statistically significant differences between the different conditions are indicated by different letters as assessed by an ANOVA test ($p < 0.05$, n=11-12 seeds per condition).

**E and F:** Polyester monomer levels in WT, *gpat457* and *myb107-2* mature Warm and Cold seeds, as indicated. Values represent means ± SD, n = 4. For each monomer, different lowercase letters indicate significant differences, as determined by Krusal-Wallis or one-way ANOVA test, $p < 0.05$

### Supplementary Figure 5

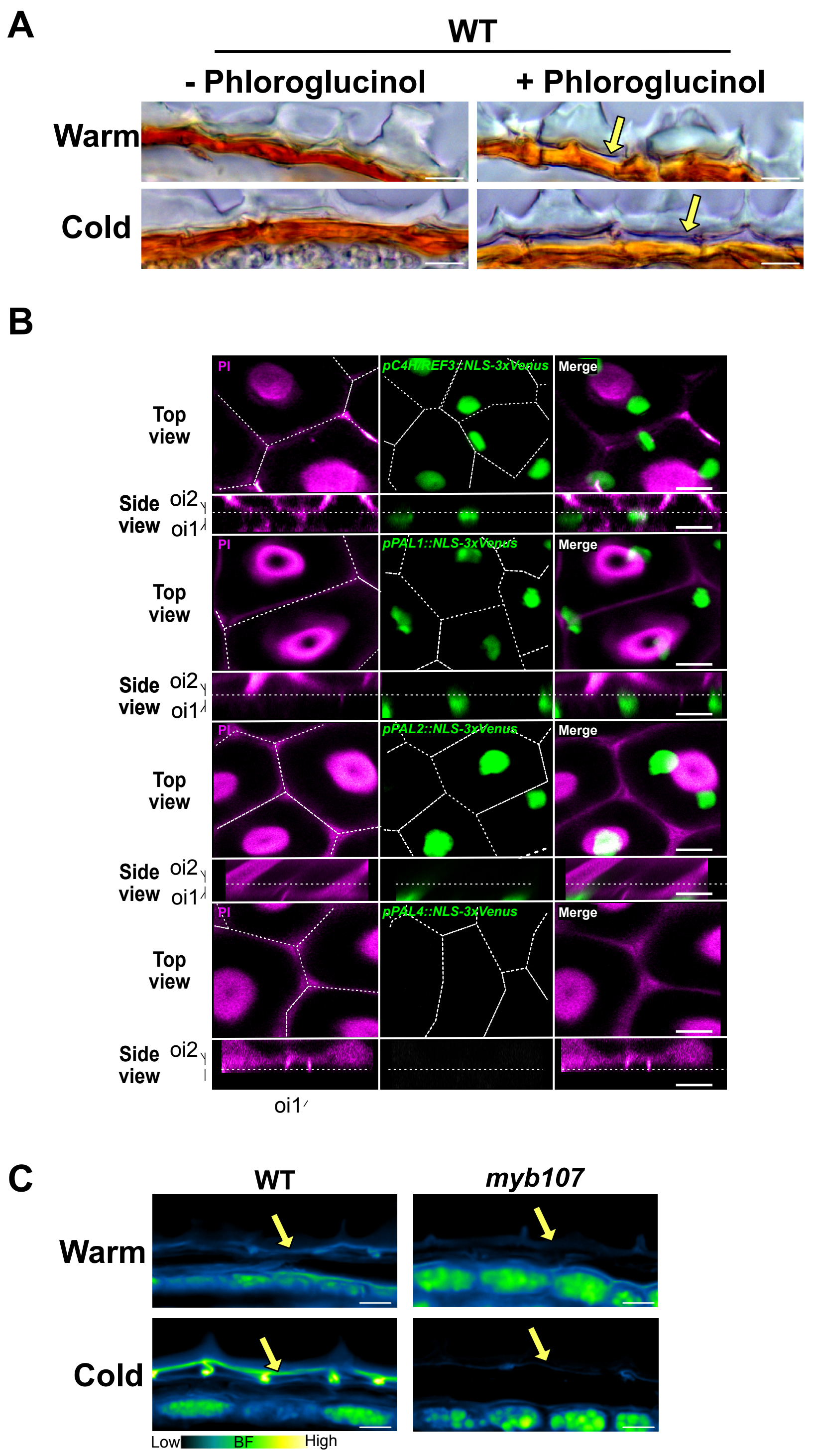


**Supplementary Figure 5: Cold promotes the polar deposition of lignin or lignin-like polymers in oi1 cells**

**A**. Pholoroglucinol staining of histological sections of WT mature Warm and Cold seeds. Yellow arrow indicates a purple signal between the oi1 and oi2 cells that is more visible in Cold seeds. Bar, 10µm.
**B**. Confocal images showing propidium iodide (PI) and Venus fluorescence in the seed mature green stage of *pC4H/REF3::NLS-3xVenus*, *pPAL1::NLS-3xVenus*, *pPAL2::NLS-3xVenus* and *pPAL4::NLS-3xVenus* transgenic plants, as indicated. Bar, 10 µm.
**C**. Basic Fuchsin (BF) staining of WT and *myb107-2* mature Warm and Cold seed sections. Yellow arrow indicates the BF signal corresponding to the polar oi1 barrier. Bar, 10µm.
